# Interactions of the *Escherichia coli* SMC-like RecN protein to different forms of DNA revealed at the single-molecule level using optical tweezers

**DOI:** 10.1101/2025.05.13.653842

**Authors:** Viktoria Dmitrievna Roshektaeva, Aleksandr Andreevich Alekseev, Alexey Dmitrievich Vedyaykin, Mikhail Alekseevich Khodorkovskii, Natalia Evgenevna Morozova

## Abstract

DNA molecule is the storage of genetic information in all living organisms. Its integrity is critical to life. However, due to the exposure to various environmental factors and endogenous agents, double-strand DNA breaks occur. Bacteria are capable to restore their genome integrity through a process called the SOS response. The key protein of SOS response is the RecA recombinase. Also critical for DNA repair is the SMC-like RecN protein, which helps RecA to find the homologous DNA template. Currently, its functions and mechanism of action remain poorly understood. In this work, using optical tweezers, we show predominant binding of RecN to ssDNA and also demonstrate a weak binding of dsDNA causing a condition similar to DNA loops formation.

## Introduction

SMC (structural maintenance of chromosomes) protein complexes are a family of ATPases that participate in the dynamic regulation of chromosome structure [1]. This class of proteins is highly conserved among prokaryotes and eukaryotes. Such complexes play a key role in chromosome structure maintenance and double-strand break repair (DSB) and are crucial for normal functioning of all types of living organisms [2]. SMC includes proteins that are the part of the eukaryotic Cohesin, Condensin, SMC5/6 complexes, and also bacterial complexes MukBEF and SMC/ScpAB [3]. The backbone of these complexes is formed by dimers of very long (more than 50 nm) coiled-coil subunits that form an ABC-like ATPase site.

In 2019, human cohesin was first demonstrated to be capable of ATP-dependent active DNA loop extrusion [4, 5], which was an important achievement and confirmed the status of SMC complexes as a new class of DNA motors. In 2020, the structures of cohesin [6, 7] and condensin [8, 9] SMC complexes were determined using cryo-electron microscopy. Analysis of these structures showed that the superhelical subunits of these complexes can undergo significant conformational changes with amplitude of more than 10 nm. Such complex conformational rearrangements are proposed to be regulated by interactions with DNA, as well as ATP binding and hydrolysis, but the mechanism of functioning of these complexes remains largely unclear.

The RecN protein studied in this work is one of the important participants in the SOS response of bacteria and also relates to the SMC. It represents independent complex, presumably formed by two protein monomers [10, 11]. Like other SMC proteins, RecN contains a long supercoiled region with an ATPase domain at the end, which upon dimerization forms an ABC-like site capable of binding two ATP molecules. It is currently known that RecN is recruited to DNA repair centers together with the RecA protein, which is the main enzyme of homologous repair and an activator of the SOS response of bacteria [12–14]. According to the *in vitro* data, RecA stimulates the DNA-dependent ATPase activity of RecN, and RecN in stimulates the process of homologous strand transfer, which is one of the key function of the RecA protein [15]. Also, it was *in vitro* found that RecN interacts with the terminal regions of single-stranded DNA and nicked regions of double-stranded DNA, and in the presence of ATP is capable of interacting with DNA molecules and forming loops from them [16]. It has been demonstrated [17] that during the SOS response in *Caulobacter*, the RecA filament undergoes translocation and remodeling under the influence of the RecN protein. Recent studies also indicate that RecN plays a critical role in enhancing the efficiency of DNA double-strand break repair by participating in processes such as RecA filament formation and homology search [18]. Our previous *in vivo* studies performed at the single-cell level, have shown that the RecN outside the SOS response is predominantly localized at the poles of the cell, and in the divisome area of dividing cells. Using *in vitro* technics including fluorescence microscopy of single DNA molecules and optical tweezers, we show that RecN predominantly binds ssDNA in an ATP-dependent manner and also has both intrinsic and single-stranded DNAstimulated ATPase activity. We also found that the RecN *E. coli*, when induced in vivo, increases the frequency of recombination exchanges by approximately 3 times [19].

Since the exact mechanism of *E. coli* RecN involvement in the double-strand break repair remains poorly understood, this work is devoted to the *in vitro* study of the nature of the interaction of RecN with single-stranded and double-stranded DNA, as well as the interaction of the RecN with different conformations of the RecA filament using single-molecule approach.

## Materials and methods

### Proteins purification

Wild-type *E. coli* RecN and RecA proteins were purified as described previously [19–21].

### DNA electrophoretic mobility shift in the presence RecN and RecA

Plasmid DNA of bacteriophage M13 was used as a DNA substrate. Four forms of DNA were used: circular single-stranded, circular double-stranded, linear single-stranded and linear double-stranded. Circular DNA was obtained by isolating plasmid DNA from *E. coli* cells infected with phage M13 using the Plasmid Miniprep kit (Evrogen). Linear DNA was obtained by restriction of the plasmid using the EcoRI restriction enzyme (Themo Scientific, Fast Digest). Partially single-stranded DNA was obtained by heating the tube at 95°C for 5 minutes and then rapidly cooling it on ice. As a result of the rapid cooling, the DNA does not anneal and remains in a partially single-stranded form.

For all binding experiments, a buffer containing 20 mM Tris-HCl, 7.7 mM KCl, 2.5 mM MgCl_2_, 1 mM DTT (pH=7.0) and 10 mM ATP (optional) was used. An excess of RecN and RecA proteins was added to the reaction, namely approximately 10^3^ protein molecules per one DNA molecule.

After mixing, the samples were incubated at 37°C for 15 minutes. Then, DNA was electrophoresed in a 1% agarose gel based on Tris-acetate buffer at a constant voltage of 110 V. After electrophoresis, the gel was incubated in TAE buffer with the addition of SYBR Gold dye (Thermo Scientific) to visualize DNA.

An analysis of the efficiency of various forms of DNA binding by the RecA and RecN proteins, as well as a mixture of RecA and RecN proteins, was carried out by the presence of a fluorescent signal from the dye in the wells and the blurring of DNA bands on the gel.

### Characterization of the RecN ATPase activity stimulation by preformed RecA filaments

To form RecA filaments, the reaction mixture containing 32 μM RecA, 96 μM poly(dT) (concentration relative to the number of bases), 2 mM ATPγS was incubated at 37°C for 20 min. Buffer R (20 mM Tris-acetate pH 7.5, 10 mM Magnesium acetate, 80 mM Potassium acetate) was used as a buffer base. After incubation, the reaction mixture with RecA filaments was purified from excess ATPγS using a Micro Bio-Spin P-30 (Bio-Rad) chromatographic spin column equilibrated with buffer R and used immediately in experiments to determine the effect of RecA filaments on the ATPase activity of RecN.

ATPase activity was registered at 37 ° C in a Buffer R with the addition of 2 mm ATP, 2 mm PEP, 30 units/ml of pyruvate kinase, 30 units/ml of LDH, 0.2 mm NADH. RecN was added in a concentration of 1 μm, pre-formed RecA filaments - in the concentration of 2 μm of RecA. The measurements were carried out using the Cary 5000 spectrophotometer (Varian). The formation of ADF during the ATP hydrolysis was chemically associated with the oxidation of NADH to NAD+ and was detected to reduce the absorption on the wavelength of 340 nm using the NADH using an extinction coefficient ε_380_=1.21 mM^−1^cm^−1^.

### The study of DNA binding properties by RecN using optical tweezers

The single-molecules experiments were performed on a double-trap optical tweezers set-up described previously [22]. Briefly the AxioImager.Z1 microscope (Carl Zeiss) equipped with a Nd:YVO4 1064 nm CW infrared laser (5 W, Spectra Physics BL-106C), a high numerical aperture oil immersion objective (100X, NA=1.25, LOMO), five-channel microfluidic flow chip (Lumicks), a high-precision piezo stage (P-561.3DD, PI) and EMCCD camera (Andor Technology, iXon Ultra 897) were used. The installation was controlled by ccustom made LabView software.

To study the dynamics of RecN-DNA interactions by the optical tweezers the linear double stranded DNA (dsDNA) molecules (11,100 bp in length) with a biotin modification of 5’- and 3’-ends of the same strand were used. Such DNA molecules were obtained as a result of restriction of the pRL574 plasmid vector with an insert of the *rpoC* gene and ligating to the sticky ends of biotinylated oligonucleotides. The experiments were carried out using optical tweezers, which allows to manipulate single DNA molecules and to apply the tension forces in the range from 0.1 to several hundreds of Piconewton (pN) and also to register the DNA length.

For DNA manipulations, polystyrene microspheres (2.1 μm, Spherotech) coated with streptavidin were used. The microspheres were held with two optical traps, and DNA was attached to microspheres through biotin-streptavidin interactions.

Combination of the optical tweezers method with a five-channel microfluidic system allowed the captured objects to be transferred between the chamber channels, thereby changing the composition of the solution around the DNA (Figure 1). The buffer solution used in single-molecule experiments was as follows: 25 mM Tris-HCl (pH 7.5), 10 mM NaCl, 5 mM MgCl_2_. The experiments were carried out at 22°C.

**Figure 1.**
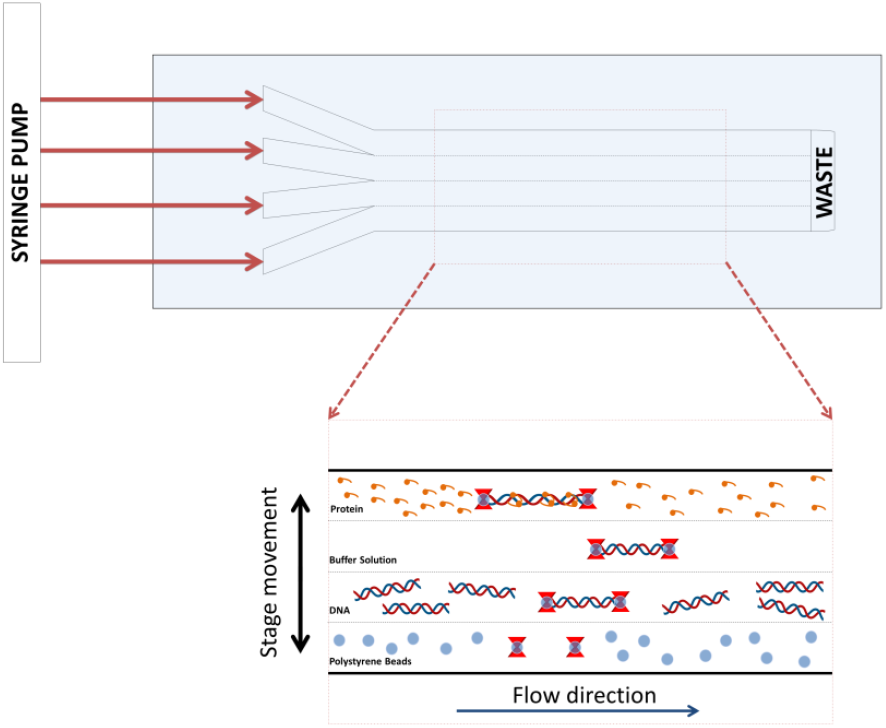
A schematic of single molecule DNA-protein binding experiment performed in microfluidic flow chamber. The first step – beads trapping; the second step – DNA capturing; the third step – DNA-protein interaction analysis

### Dynamics of RecN interaction with dsDNA

To determine the dynamics of the interaction of RecN with dsDNA, the dependences of tension on DNA length were measured during stretching and relaxation cycles after incubation of the double-stranded DNA molecule in the presence of 500 nM RecN in the relaxed state.

### Dynamics of RecN interaction with ssDNA

To conduct experiments with single-stranded DNA (ssDNA), the ssDNA molecules were obtained from dsDNA by force melting method directly during the experiment. Force melting method represents mechanical overstretching of dsDNA with a force of more than 80 pN, leading to the distruction of hydrogen bonds between complementary DNA strands. Due to the attachment of DNA to the microspheres through the biotin-modified 5’- and 3’-ends of the same DNA strand in the duplex molecule, the melting of dsDNA caused by mechanical overstretching leads to the fact that one DNA strand remains fixed between the microspheres, and the complementary strand is irreversibly distanced due to diffusion and the viscous friction force from the flow. The complete transition of DNA from the double-stranded to the single-stranded form can be tracked by measuring the DNA stretching curves before and after melting.

### Study of RecN interaction with different RecA filament conformations using optical tweezers

The active RecA filament formation on ssDNA was carried out using the optical tweezers. The filament was formed in the presence of 1 μM RecA, 3 mM ATP, 1 mM PEP, 10 U/ml pyruvate kinase. The formation of the RecA filament on ssDNA leads to its length increase which could be monitored in the experiment. The RecN protein was added at a concentration of 500 nM. The experiments were carried out at 22°C in a buffer containing 25 mM Tris-HCl (pH 7.5), 10 mM NaCl, 5 mM MgCl2. To obtain an inactive RecA-ssDNA filament, a RecA-ssDNA filament preformed in the presence of ATP was transferred to the channel of the chamber with a buffer without ATP, as a result of which the RecA-ssDNA filament reversibly passed into an inactive conformation as a result of total hydrolysis of ATP molecules bound in the ATPase sites of the filament.

## Results

### Binding of DNA by RecA and RecN simultaneously causes a shift of different forms of DNA in an agarose gel, and this shift is significantly higher than that which occurs after the interaction of each of these proteins with DNA

Analysis of the electrophoretic mobility of DNA in the presence of RecN and RecA proteins in 1% agarose gel showed that RecN itself has DNA-binding activity, which is more pronounced on linear ssDNA (Figure 2A). RecA also has DNA-binding activity, which is more pronounced on linear dsDNA and linear ssDNA (Figure 2B). However, the addition of RecA and RecN to the binding reaction simultaneously significantly improves the binding of all forms of DNA (Figure 2A and 2B). To check this result we perform the estimation of the RecN ATPase activity stimulation by preformed RecA filaments.

**Figure 2.**
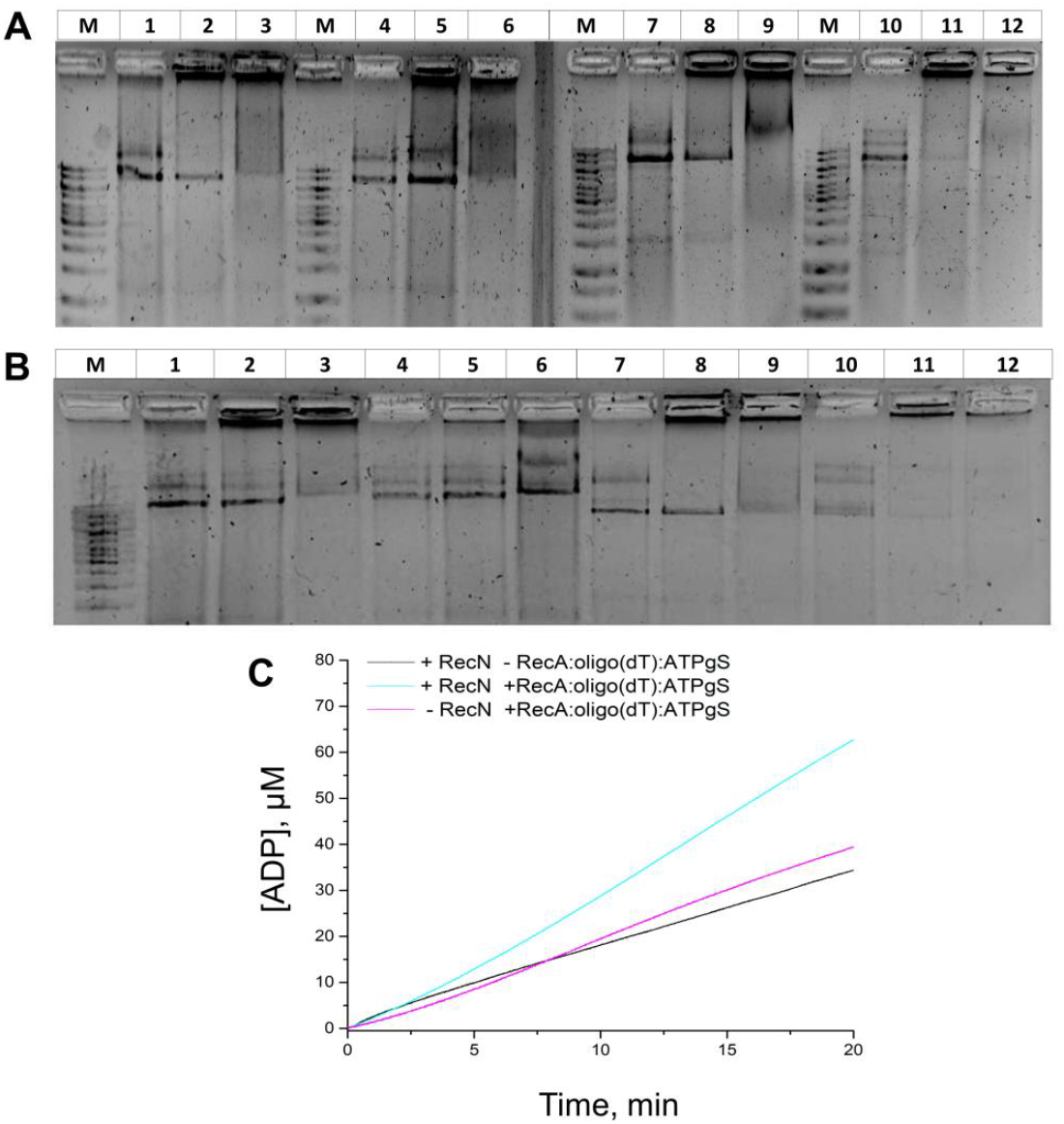
RecA and RecN DNA binding properties. **A** – Analysis of electrophoretic mobility of DNA in the presence of RecN and RecA proteins. DNA electrophoresis was performed in 1% agarose gel. **M** – DNA Ladder 1kb; **1** – circular dsDNA; **2** – circular dsDNA, RecN, 10 mM ATP; **3** – circular dsDNA, RecN, RecA, 10 mM ATP; **4** – circular ssDNA; **5** – circular ssDNA, RecN, 10 mM ATP; **6** – circular ssDNA, RecN, RecA, 10 mM ATP; **7** – linear dsDNA; **8** – linear dsDNA, RecN, 10 mM ATP; **9** – linear dsDNA, RecN, RecA, 10 mM ATP; **10** – linear ssDNA; **11** – linear ssDNA, RecN, 10mM ATP; **12** – linear ssDNA, RecN, RecA, 10mM ATP. RecN itself has DNA-binding activity, which is most clearly expressed on linear ssDNA. Adding the RecA protein to the reaction significantly improves the binding of all forms of DNA; **B** – Analysis of DNA electrophoretic mobility in the presence of RecA and RecN proteins. DNA electrophoresis was performed in 1% agarose gel. **M** – DNA Ladder 1kb; **1** – circular dsDNA; **2** – circular dsDNA, RecA, 10 mM ATP; **3** – circular dsDNA, RecN, RecA, 10 mM ATP; **4** – circular ssDNA; **5** – circular ssDNA, RecA, 10 mM ATP; **6** – circular ssDNA, RecN, RecA, 10 mM ATP; **7** – linear dsDNA; **8** – linear dsDNA, RecA, 10 mM ATP; **9** – linear dsDNA, RecN, RecA, 10 mM ATP; **10** – linear ssDNA; **11** – linear ssDNA, RecA, 10mM ATP; **12** – linear ssDNA, RecN, RecA, 10mM ATP. RecA itself has DNA-binding activity, predominantly expressed on linear dsDNA and linear ssDNA. However, the addition of RecN protein to the reaction significantly improves the binding of all forms of DNA; **C** – RecN ATPase activity registration in the absence and presence of preformed RecA filaments

### The increase in the ATP hydrolysis rate when preformed RecA filaments are added to RecN is an additive effect

The results of RecN ATPase activity measurements in the absence and presence of preformed RecA filaments, as well as the ATPase activity of preformed RecA filaments in the absence of RecN are shown in the Figure 2C. When preformed RecA filaments were added to the RecN protein, the observed ATP hydrolysis rate was higher than in the case when RecA filaments were not added. However, during control measurements, a significant level of ATPase was detected in RecA filaments pre-formed in the presence of ATPgS in the reaction without RecN. This effect is due to the fact that ATPgS is a slowly hydrolyzed analogue of ATP. Because of this, partial replacement of nucleotide cofactor molecules bound in the

ATPase sites of the filament can occur in the presence of ATP. Analysis of the obtained data indicates that an increase in the ATP hydrolysis rate, when adding pre-formed RecA filaments to RecN, is an additive effect. Stimulation of the ATPase activity of *E. coli* RecN by *E. coli* RecA filaments under described conditions is not observed.

### Single molecule studies reveal that RecN interacts with double-stranded DNA and causes its compaction

To determine the dynamics of RecN interaction with dsDNA, the dependences of tension on DNA length were measured during stretching and relaxing cycles after incubating a double-stranded DNA molecule in a relaxed state in the presence of 500 nM RecN.

The stretching curves of unbound dsDNA can be described by the Worm-like-chain model, the parameters of which are the contour and persistent lengths of the polymer. At an end-to-end distance (the length observed in the experiment) significantly smaller than the contour length of DNA, the recorded tension is close to zero. As the distance between the microspheres gets closer to the contour length of DNA, the value of the recorded tension force begins to increase (Figure 3A).

**Figure 3.**
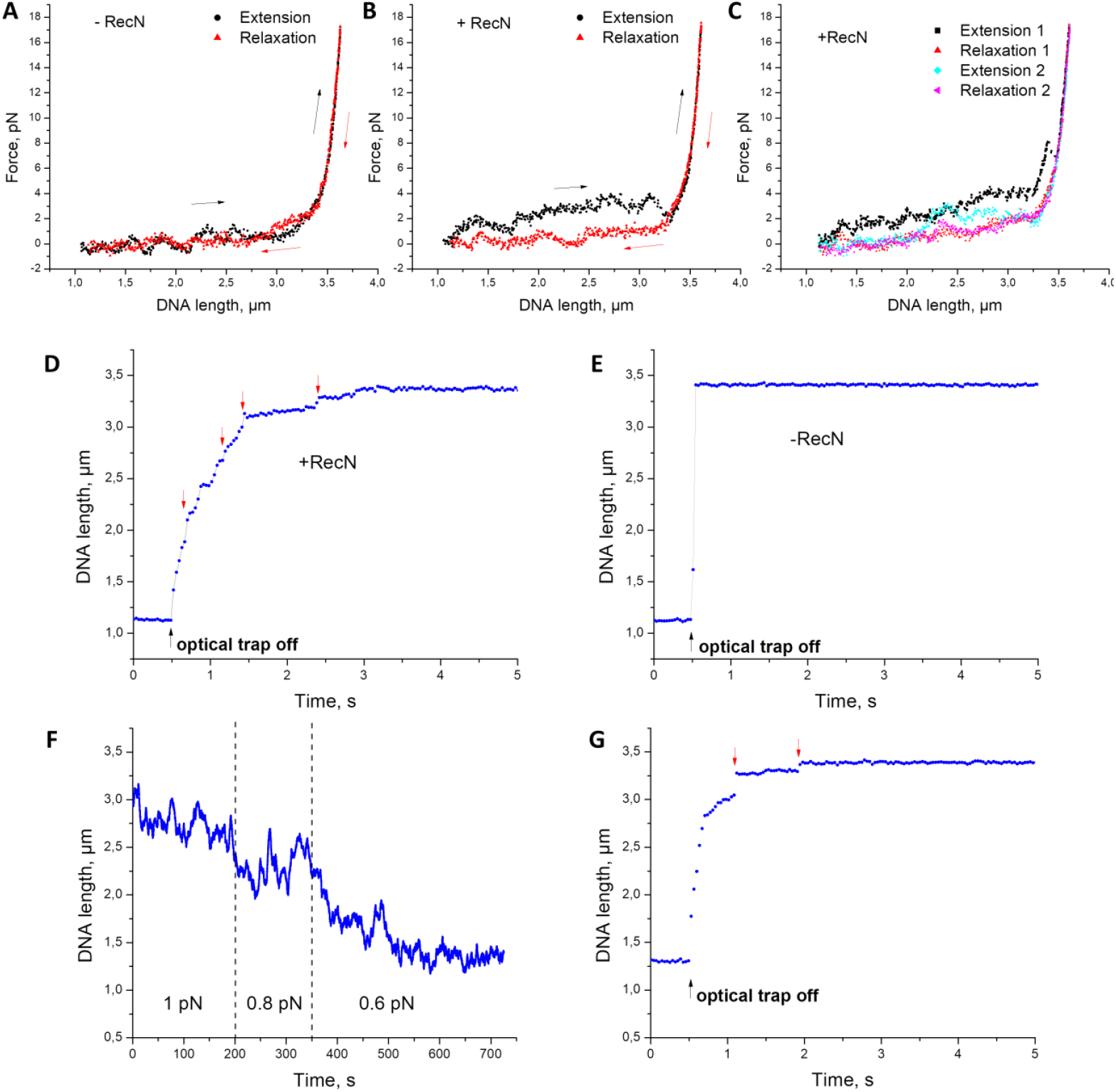
Interaction of RecN with dsDNA. **A** – Extension-relaxation curves of unbound dsDNA; **B** – Extension-relaxation curves of dsDNA in the channel with 500 nM RecN after incubation in the relaxed state for 100 seconds; **C** – Repetition of stretching and relaxation cycles of dsDNA after incubation of dsDNA in a channel with 500 nM RecN; **D** – Dynamics of dsDNA length change caused by its straightening due to the action of a constant force (~3 pN) from the flow after incubation in channel with 500 nM RecN; **E** – Dynamics of free dsDNA length change caused by its straightening due to the action of a constant force (~3 pN) from the flow; **F** – Compaction dynamics of dsDNA during incubation in a channel with 500 nM RecN under the application of a constant tension force; **G** – Dynamics of dsDNA length change caused by its straightening due to the action of a constant force (~3 pN) from the flow after incubation in the channel with RecN in accordance with **F**

Incubation of dsDNA with RecN led to an increase in the tension recorded during extension of the DNA-protein complex in the region of small lengths by 1–3 pN (Figure 3B), which may be a consequence of the formation of intramolecular bridges during the binding of RecN to DNA and their subsequent destruction under the action of mechanical force.

The observed effect was most pronounced during incubation of dsDNA in relaxed state in the presence of RecN. To reach relaxed state of used DNA molecules (the length about 3.7 μm) the distance between the microspheres during incubation was approximately 1 μm. An increase in the incubation time led to an increase in the difference between the stretching and relaxation curve. In case when stretching and relaxation cycles was repeated without additional incubation of DNA in a relaxed state, the difference between the stretching and relaxation curve became smaller (Figure 3C), which indicates that the restoration of the structures formed by the RecN protein on dsDNA after their mechanical stretching does not occur immediately.

The obtained data indicate that the structures formed by the RecN protein on dsDNA are degraded when a tension in the range of 2-4 pN is applied. Due this, we record the dynamics of dsDNA length changes at a constant tension force after incubation with RecN in a relaxed state. For this purpose, after incubation of dsDNA in RecN, one of the optical traps was switched off using an optical shutter, as a result of which a constant force of about 3 pN was applied to the free microsphere by the flow, which led to DNA straightening under the action of this force (Figure 3D, 3E). Straightening of dsDNA after incubation in a relaxed state in the channel with RecN occurred with a delay. Also, in the obtained time dependences of DNA length changes, multiple sharp increases in length are observed (marked in Figure 3D with red arrows). Such behavior may indicate the destruction of individual intramolecular bridges formed by the RecN protein.

We also record the dynamics of changes in the dsDNA length during incubation in 500 nM RecN under the application of a small constant tension force (1; 0.8; 0.6 pN). In this case, the constant force applied to the DNA was maintained at the hardware level by shifting the movable trap in the feedback loop mode. As a result, it was found that RecN gradually compacts the dsDNA (Figure 3F) under a tension of up to 1 pN. The compaction of dsDNA by the RecN protein under such conditions was additionally confirmed by recording the dynamics of changes in the dsDNA length when one of the optical traps was switched off and the dsDNA was straightened due to the action of a constant force of about 3 pN from the flow (Figure 3G).

### Single molecule studies reveal that RecN binds single-stranded DNA and such binding is relatively strong

Incubation of RecN with ssDNA resulted in pronounced changes in the ssDNA stretching curve (Figure 4A), which, as in the case of dsDNA, can be characterized as DNA compaction upon binding to RecN. After incubation in the presence of RecN, the length of the DNA-protein complex was shorter at the same tension force than the length of unbound ssDNA. Hysteresis was observed in the extension-relaxation curves of the RecN-ssDNA complex, indicating the formation of structures by the RecN protein that undergoes changes upon application of mechanical force: at the same distances between the microspheres, the tension recorded during stretching was greater than during relaxation. Increasing the incubation time of ssDNA in RecN resulted in a greater degree of ssDNA compaction as can be seen from the graphs with stretch-relaxation curves in Figure 4B. The changes in the ssDNA stretch curve were also maintained when ssDNA was transferred from the RecN-containing channel to the buffered channel without free RecN, indicating a high affinity of RecN for ssDNA (Figure 4C).

**Figure 4.**
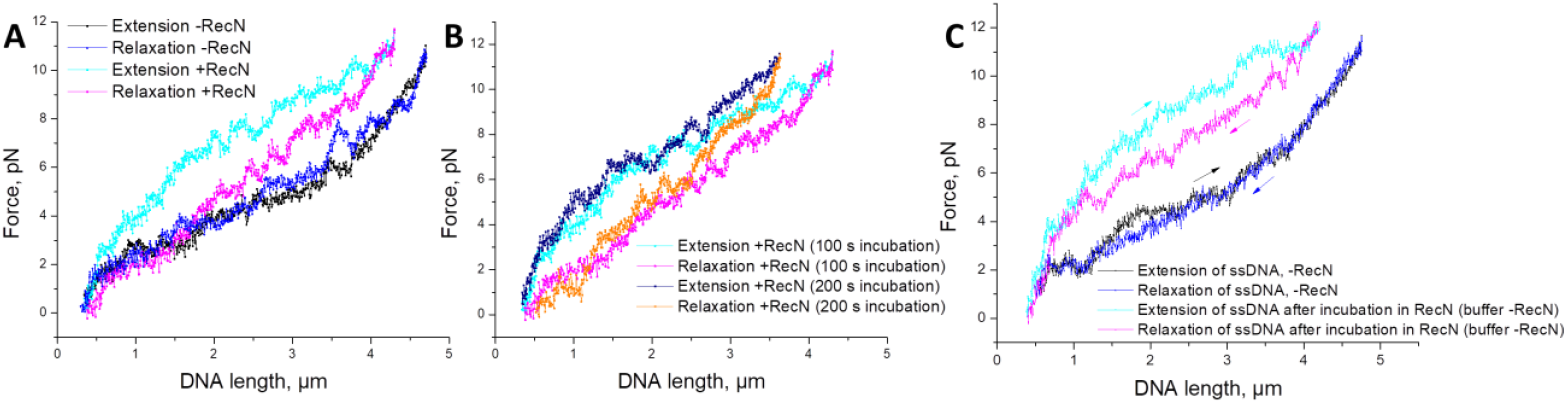
Interaction of RecN with ssDNA. **A** – Extension-relaxation curves of free ssDNA and ssDNA after incubation in 500 nM RecN for 100 seconds; **B** – Extension-relaxation curves of free ssDNA and ssDNA after incubation in 500 nM RecN for 100 seconds and 200 seconds; **C** – RecN remains bound to ssDNA in buffer without free RecN. Black and blue show the stretch-relaxation curves of unbound ssDNA. Cyan and magenta show the extension-relaxation curves in buffer without free RecN after incubation of ssDNA with 500 nM RecN

Thus here we have shown that binding of ssDNA by RecN is stronger, that in the case of dsDNA and the protein remains bound to ssDNA also in absence of free RecN.

### Interactions of RecN with different conformations of RecA filament

In bacterial cells, the RecA filament formed on ssDNA participates in recombination repair and triggers the SOS response mechanism in response to significant DNA damage and stress caused by antibacterial drugs. It is known that the RecA-ssDNA filament can be in an active or inactive conformation. The active conformation is realized in the presence of ATP, and the inactive conformation is realized in the presence of ADP or in the absence of a nucleotide cofactor. The inactive conformation of the RecA filament is characterized by a smaller helix pitch compared to the active conformation. It is also known that regions of inactive conformation appear in the structure of the active RecA-ssDNA filament located in a medium with ATP, as a result of a local ATP hydrolysis and serve as a substrate for binding by some proteins – regulators of RecA activity.

We studied the interactions of RecN with the active and inactive conformations of the RecA-ssDNA filament using the optical tweezers method, and also studied the dynamics of the active RecA-ssDNA filament formation in the presence of RecN.

According the results, described above, the RecN protein has a strong affinity for ssDNA and is able to have a compacting effect on it. It could potentially lead to competition between RecA and RecN for binding to ssDNA and obstruct the formation of the RecA-ssDNA filament. In the current study, we showed that the presence of RecN together with RecA does not interfere with the formation of the RecA filament on ssDNA. The dynamics of ssDNA length changes during incubation in the presence of RecA and ATP as well as in the presence of RecA, ATP and RecN were identical (Figure 5A).

**Figure 5.**
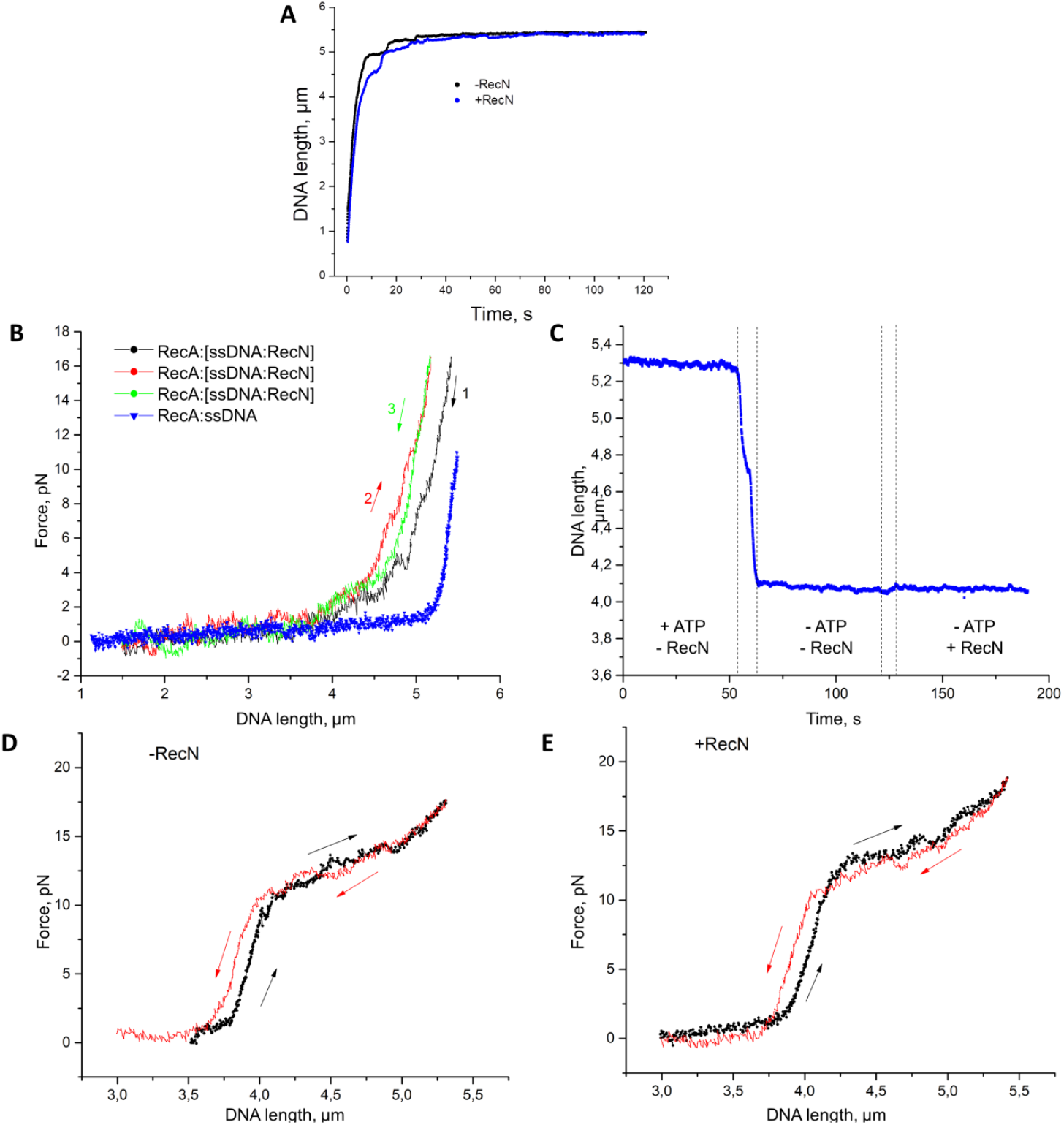
Interactions of RecN with different conformations of the RecA filament. **A** – Dynamics of ssDNA length changes during the formation of the RecA-ssDNA filament under the application of a constant tension force of 12 pN in the absence and presence of RecN; **B** – Extension curves of the active RecA-ssDNA filament. Black, red and green colors show successively recorded relaxation and extension curves of the RecA filament formed on ssDNA after preliminary incubation of ssDNA with RecN. Blue color shows the extension curve of the RecA filament formed without preliminary incubation of ssDNA with RecN; **C** – Transition of the RecA-ssDNA filament to an inactive conformation and the dynamics of the inactive RecA-ssDNA filament length in the absence and presence of RecN. A constant tension force of 3 pN was applied to the filament during the experiment; **D, E** – Extension - relaxation curves of the inactive RecA-ssDNA filament in the absence (**D**) and presence of RecN (**E**)

At the same time, it was found that in the case of preliminary incubation of ssDNA with RecN and without RecA, the mechanical properties of the RecA-ssDNA filament formed afterwards were somewhat different from the case when the RecA-ssDNA filament was formed without preliminary incubation of ssDNA in RecN (Figure 5B). Preliminary incubation of ssDNA with RecN led to the fact that the RecA filament subsequently formed on such ssDNA had a shorter length and was less rigid.

Also, to test the effect of RecN on the active conformation of the RecA-ssDNA filament, the dynamics of the formed filament length change were measured during incubation without and with RecN under conditions of a constant tension force of 3 pN. The length of the RecA-ssDNA filament at a force of 3 pN remained almost stable both in the absence and in the presence of RecN. Measurements of the extension and relaxation curves did not reveal the compaction effects caused by the RecN protein on the RecA-ssDNA filament, similar to the effects observed on ssDNA and dsDNA. To determine the effect of RecN on the inactive RecA-ssDNA filament, the dynamics of the change in the filament length were measured during incubation in the buffer without RecN and in the buffer with RecN under a constant tension force of 3 pN (Figure 5C). No effect of RecN on the inactive RecA-ssDNA filament length dynamics was detected. Moreover, the extension-relaxation curves of the inactive RecA-ssDNA filament obtained in the absence and presence of RecN were identical (Figure 5D, 5E).

## Discussion

Analysis of the electrophoretic mobility of DNA in the presence of RecN and RecA proteins in 1% agarose gel showed that RecN itself has DNA-binding activity, most clearly expressed on linear ssDNA. And RecA itself also has DNA-binding activity, most pronounced on linear dsDNA and linear ssDNA. However, presence of both RecA and RecN proteins significantly improves the binding of all forms of DNA.

However, the analysis of RecN ATPase activity in the absence and presence of preformed RecA filament show that there is no significant synergistic effect of these two proteins is observed. The observed increase in ATPase activity is the sum of the RecA and RecN ATPase activities and represents an additive effect.

Single-molecules approach allowed to measure changes in the mechanical properties of DNA substrates upon interaction with RecN, visualize the effect of dsDNA compaction and to estimate the magnitude of the compression force at about 1 pN. It was also possible to register the increasing dynamics of the compacted dsDNA at application of the pulsed force of about 3 pN in the opposite to compression force direction - DNA length increased up to contour value within 2-3 seconds. The observed effect of such essential decreasing of the dsDNA apparent length at such low compaction forces may be due to formation of loop extrusion created by RecN extrusion complex.

In the case of ssDNA, the changes in the tension-length dependence upon interaction with RecN are more pronounced. The application force of 3-4 pN is insufficient to eliminate the compaction of ssDNA caused by RecN binding. At the same time, the binding of RecN to ssDNA is preserved upon transfer of the complex to a solution without free RecN, which indicates a high affinity of RecN for DNA in the single-stranded form. This was also confirmed in our previous study [19].

The study of RecN interactions with various conformations of the RecA filament showed that no special effect on the conformational states of the RecA filament is observed. In this case, the filament assembles on ssDNA in the presence of RecA alone, and in the presence of RecA and RecN in the same way.

However, if the ssDNA molecule is pre-incubated in RecN, the dynamics of the RecA filament assembly changes, which may indicate the presence of competition between RecA and RecN for binding to ssDNA. The filament built in the presence of RecA and RecN is more stable and loses less of its length during relaxation. The obtained results indicate the absence of direct interaction of RecN with the RecA-ssDNA filament in the active and inactive conformations. At the same time, the regulation of the RecA-ssDNA filament activity by the RecN protein can be carried out through the interaction of RecN with DNA.

## Acknowledgements

The research was supported by Russian Science Foundation, project No. 24-74-00085, https://rscf.ru/project/24-74-00085/ to NEM and project No. 24-74-10022, https://rscf.ru/project/24-74-10022/ to ADV. The work was carried out using scientific equipment of the Center of Shared Usage “The analytical center of nano and biotechnologies of SPbPU”.

## References

1. Reyes, E.D., et al., RecN is a cohesin-like protein that stimulates intermolecular DNA interactions in vitro. J Biol Chem, 2010. 285(22): p. 16521–9.

2. Kim, E., R. Barth, and C. Dekker, Looping the Genome with SMC Complexes. Annu Rev Biochem, 2023. 92: p. 15–41.

3. Hirano, T., At the heart of the chromosome: SMC proteins in action. Nat Rev Mol Cell Biol, 2006. 7(5): p. 311–22.

4. Davidson, I.F., et al., DNA loop extrusion by human cohesin. Science, 2019. 366(6471): p. 1338–1345.

5. Kim, Y., et al., Human cohesin compacts DNA by loop extrusion. Science, 2019. 366(6471): p. 1345–1349.

6. Shi, Z., et al., Cryo-EM structure of the human cohesin-NIPBL-DNA complex. Science, 2020. 368(6498): p. 1454–1459.

7. Higashi, T.L., et al., A Structure-Based Mechanism for DNA Entry into the Cohesin Ring. Mol Cell, 2020. 79(6): p. 917–933 e9.

8. Lee, B.G., et al., Cryo-EM structures of holo condensin reveal a subunit flip-flop mechanism. Nat Struct Mol Biol, 2020. 27(8): p. 743–751.

9. Kong, M., et al., Human Condensin I and II Drive Extensive ATP-Dependent Compaction of Nucleosome-Bound DNA. Mol Cell, 2020. 79(1): p. 99–114 e9.

10. McLean, E.K., J.S. Lenhart, and L.A. Simmons, RecA Is Required for the Assembly of RecN into DNA Repair Complexes on the Nucleoid. J Bacteriol, 2021. 203(20): p. e0024021.

11. Pellegrino, S., et al., Structural and functional characterization of an SMC-like protein RecN: new insights into double-strand break repair. Structure, 2012. 20(12): p. 2076–89.

12. Sanchez, H., et al., Recruitment of Bacillus subtilis RecN to DNA double-strand breaks in the absence of DNA end processing. J Bacteriol, 2006. 188(2): p. 353–60.

13. Keyamura, K., et al., RecA protein recruits structural maintenance of chromosomes (SMC)-like RecN protein to DNA double-strand breaks. J Biol Chem, 2013. 288(41): p. 29229–37.

14. Lesterlin, C., et al., RecA bundles mediate homology pairing between distant sisters during DNA break repair. Nature, 2014. 506(7487): p. 249–53.

15. Uranga, L.A., et al., The cohesin-like RecN protein stimulates RecA-mediated recombinational repair of DNA double-strand breaks. Nat Commun, 2017. 8: p. 15282.

16. Sanchez, H., et al., Dynamic structures of Bacillus subtilis RecN-DNA complexes. Nucleic Acids Res, 2008. 36(1): p. 110–20.

17. Chimthanawala, A., et al., SMC protein RecN drives RecA filament translocation for in vivo homology search. Proc Natl Acad Sci U S A, 2022. 119(46): p. e2209304119.

18. Noda, S., et al., RecN spatially and temporally controls RecA-mediated repair of DNA double-strand breaks. J Biol Chem, 2023. 299(12): p. 105466.

19. Roshektaeva, V.D., et al., Features of the DNA Escherichia coli RecN interaction revealed by fluorescence microscopy and single-molecule methods. Biochem Biophys Res Commun, 2024. 716: p. 150009.

20. Keyamura, K. and T. Hishida, Topological DNA-binding of structural maintenance of chromosomes-like RecN promotes DNA double-strand break repair in Escherichia coli. Commun Biol, 2019. 2: p. 413.

21. Drees, J.C., S.L. Lusetti, and M.M. Cox, Inhibition of RecA protein by the Escherichia coli RecX protein: modulation by the RecA C terminus and filament functional state. J Biol Chem, 2004. 279(51): p. 52991–7.

22. Pobegalov, G., et al., Deinococcus radiodurans RecA nucleoprotein filaments characterized at the single-molecule level with optical tweezers. Biochem Biophys Res Commun, 2015. 466(3): p. 426–30.

